# Standardizing protein corona characterization in nanomedicine: a multi-center study to enhance reproducibility and data homogeneity

**DOI:** 10.1101/2024.04.25.591109

**Authors:** Ali Akbar Ashkarran, Hassan Gharibi, Seyed Majed Modaresi, Amir Ata Saei, Morteza Mahmoudi

## Abstract

Our recent findings reveal substantial variability in the characterization of identical protein corona across different proteomics facilities, demonstrating that protein corona datasets are not easily comparable between independent studies. We have shown that heterogeneity in the final composition of the identical protein corona mainly originates from variations in sample preparation protocols, liquid chromatography mass spectrometry (LC-MS) workflows, and raw data processing. Here, to address this issue, we developed standardized protocols and unified sample preparation workflows, and distributed identical protein corona digests to several proteomics centers that performed better in our previous study. Additionally, we examined the influence of using similar mass spectrometry instruments on data homogeneity. Furthermore, we evaluated whether standardizing database search parameters and data processing workflows could enhance data uniformity. More specifically, our new findings reveal a remarkable, stepwise improvement in protein corona data consistency across various proteomics facilities. Streamlining the whole workflow results in a dramatic increase in protein ID overlaps from 11% for good centers to 40% across core facilities that utilized similar instruments and were subjected to a uniform database search. This comprehensive analysis identifies key factors contributing to data heterogeneity in mass spectrometry-based proteomics of protein corona and plasma-related samples. By streamlining these processes, our findings significantly advance the potential for consistent and reliable nanomedicine-based diagnostics and therapeutics across different studies.

## Introduction

The protein/biomolecular corona is a dynamic biological layer that develops on the surface of nanoparticles when they are exposed to biological fluids including plasma.^1, 2, 3, 4^ This biological layer transforms nanoparticles by endowing them with a new identity that influences how they are recognized and interact within biological systems, thus defining their safety, biodistribution, diagnostic and therapeutic efficacy.^3, 5, 6^ Despite significant advances in nanomedicine, a limited understanding of nanoparticles’ biological identity remains a major barrier to their successful clinical translation.^7^ A profound comprehension of the protein corona composition is crucial for predicting their *in vivo* biological fate, which is essential for their clinical application and the advancement of future nanomedicine technologies.^7^

Over the past decade, extensive research has aimed to enhance the reproducibility of nanobio interfaces.^8, 9^ Yet, the critical role of liquid chromatography mass spectrometry (LC-MS) in ensuring the robustness and reproducibility of protein corona data—a pivotal method for identifying and quantifying protein corona composition^10, 11, 12^—has received insufficient attention.^10, 11, 12, 13^ Our recent studies show significant discrepancies in identical protein corona outcomes and interpretations across various proteomics facilities, attributed to differences in sample preparation protocols, LC-MS workflows, instrumentation, and data processing.^13^ Consequently, there is an urgent need to develop a standardized protocol for protein corona analysis to reduce data heterogeneity and facilitate comparability across studies.^13, 14^

Our previous research indicates that major disparities in the characterization of the protein corona composition are primarily due to variations in sample preparation protocols, LC-MS workflows, and data processing.^13^ We also revealed that harmonizing database search and data processing can significantly reduce the observed heterogeneity among core facilities.^15^ Different proteomics core facilities employ a range of sample preparation methods, instrumentation, quantification approaches, search parameters, and data processing techniques. These differences can significantly bias the outcomes of protein corona analyses of identical samples. Furthermore, the multistep nature of sample preparation—which includes protein denaturation, reduction, alkylation, digestion, and desalting/cleaning—introduces additional variability due to the diverse chemicals and reagents used across different laboratories.^16, 17, 18^ Additionally, the use of various LC and MS systems contributes to another layer of variation and heterogeneity that affects the final characterization of the protein corona.^19, 20, 21^

The primary goal of this follow-up study is to minimize the main sources of variations and improve the protein corona data homogeneity, focusing on harmonizing sample preparation protocols, LC-MS instrumentation and workflows, database search, and processing strategies. Ultimately, similar to other nanomedicine techniques and methods^6^, we aim to establish a standardized protocol that will streamline the analysis of the protein corona across different core facilities, facilitating the comparability of protein corona datasets from various studies. More specifically, here, we prepared identical ready-to-inject batches of digested protein corona samples and sent them to various proteomics centers for analysis. We used an identical on-bead digestion protocol for all protein corona samples and submitted the final 4 (out of 6) identical batches of dried peptides to 4 proteomics core facilities in the United States that had provided relatively higher quality data in on our previous study (called good centers) based on protein and peptide counts, coefficients of variation (CV) of technical replicates, and the median sequence coverage.^13^ In addition, to probe the role of LC-MS instrument in heterogeneity of protein corona data, we also sent the another 4 identical batches of dried peptides to 4 proteomics core facilities that had similar instrumentation and were expected to perform similarly (i.e., identical LC and similar MS types; called similar centers). Two of these centers were shared with the above (good) centers, so in total we sent identical samples to 6 core facilities. Finally, we also performed a uniform database search on the raw data retrieved from the 4 centers with similar instruments, where the sample preparation protocol, instrumentation, and data processing would all be streamlined. In summary, we conducted the study in 3 steps: 1) unifying the sample preparation protocol; 2) unifying sample preparation protocol and the instrumentation used, and 3) unifying sample preparation protocol, instrumentation, and data processing. We show that implementation of each step reduces data heterogeneity of the protein corona composition across core facilities which can be used for analysis of protein corona in the future, enabling rapid developments in nanomedicine-based diagnostics and therapeutics.

## Results

The overall workflow of the present study is illustrated in **Fig. 1**. We employed commercially available plain polystyrene nanoparticles with an average diameter of 78.8 nm^13^. Consistent with our previously published protocols, all protein corona-coated nanoparticles were prepared under identical conditions (refer to reference^13^ for details). **Supplementary Fig. 1** illustrates the dynamic light scattering (DLS), zeta potential, and transmission electron microscopy (TEM) analyses of both bare and protein corona-coated nanoparticles. Notably, our prior research indicated minimal batch-to-batch variation in the physicochemical properties of the protein corona-coated nanoparticles. Bare nanoparticles were monodispersed with a narrow size distribution, measuring an average size of 78.8 ± 0.0 nm and a surface charge of -30.8 ± 0.8 mV. After exposure to human plasma, the average size increased to 111.3 ± 9.6 nm, and the surface charge changed to -10.2 ± 0.4 mV, confirming the formation of the protein corona. Detailed TEM analysis further elucidated the changes in size and morphology before and after protein corona formation, showing a distinct dark layer indicating the presence of the protein layer on the nanoparticles (**Supplementary Fig. 1e**).^22, 23, 24^ The protein concentration in each batch, approximately 1.6 μg, was quantified using a bicinchoninic acid assay (BCA) to ensure proper sample preparation for LC-MS analysis. These fully characterized samples were subsequently digested using a standardized protocol as described in the experimental section and dispatched to various proteomics facilities for analysis.

**Fig. 1.**
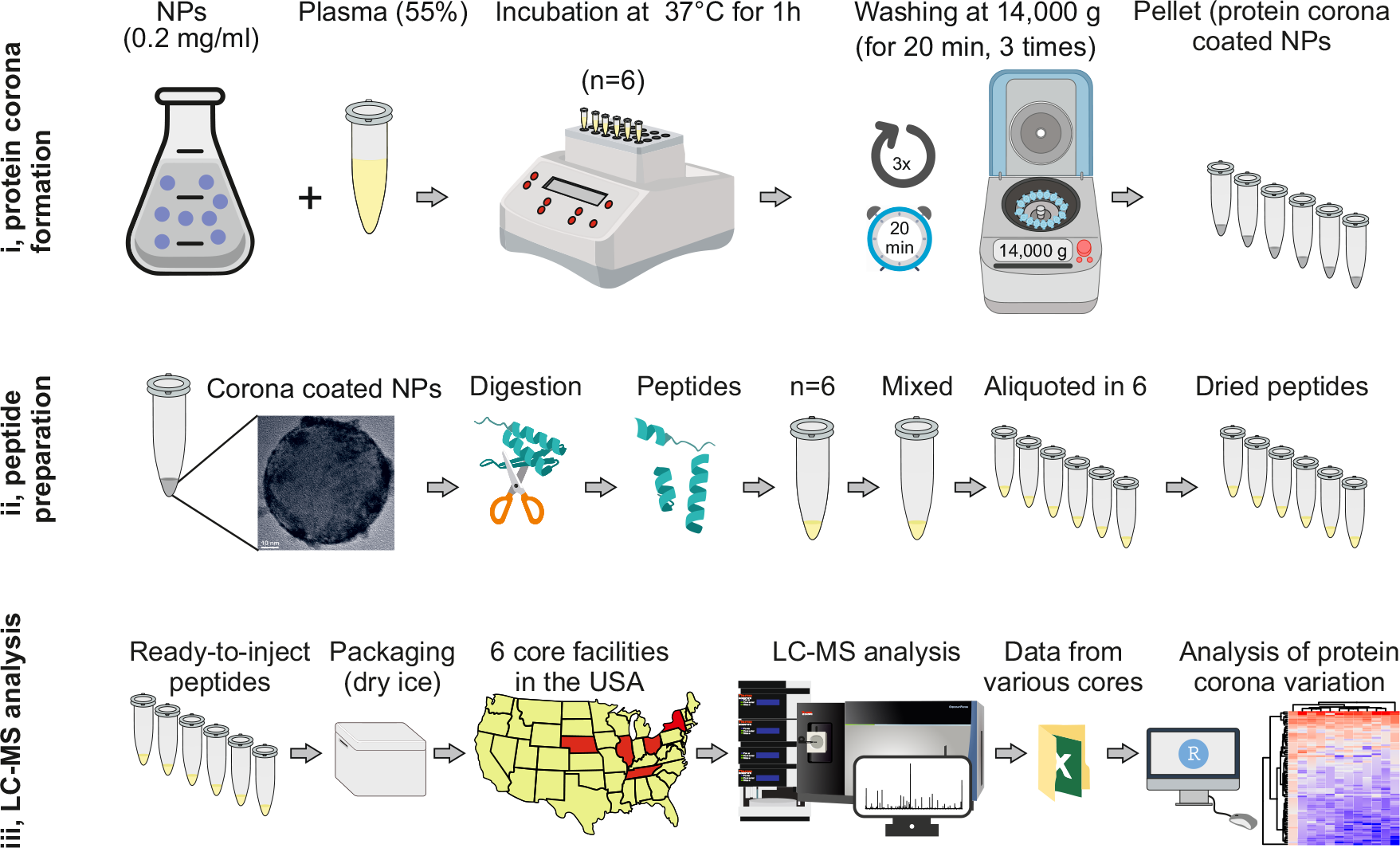
Schematic showing overall workflow of the study. After formation of 6 similar batches of protein corona–coated polystyrene nanoparticles, each individual batch was fully digested to peptides using our in-house developed protocol, mixed and aliquoted and dried. The resulting identical aliquots were shipped to 6 different proteomics core facilities across the USA (one state includes two proteomics center) to investigate the homogeneity/heterogeneity of the protein corona composition on the surface of the nanoparticles. Out of these 6 core facilities, 2 were designated as both good and similar. The cryo-TEM image of the corona-coated nanoparticle is reproduced here with permission from reference^13^.

### The impact of unifying sample preparation protocol and instrumentation

We submitted 4 identical batches of dried peptides to 4 core facilities that had better performance in terms of protein counts, peptide counts, coefficients of variation (CV) of technical replicates, as well as the median sequence coverage based on our previous findings^13^ (see **Supplementary Information** for details regarding each core facility and the associated protocols). Furthermore, 4 identical batches were submitted to the 4 proteomics core facilities in the USA with similar instrumentation (with regards to LC and MS types)^13^. These centers used the same LC system (Dionex Ultimate 3000) and MS systems that should largely produce similar results (two used Fusion Lumos, one Fusion and one HF-X system). All centers were asked to analyze the samples over a 120 min gradient using label-free quantification (LFQ). Since good performance in our original study was also taken into account ^13^, we could not identify 4 core facilities with exact same MS instruments and had to select these centers for analysis. The LC type and MS instruments as well as good/similar desginations are summarized in **Table 1**.

**Table 1.**
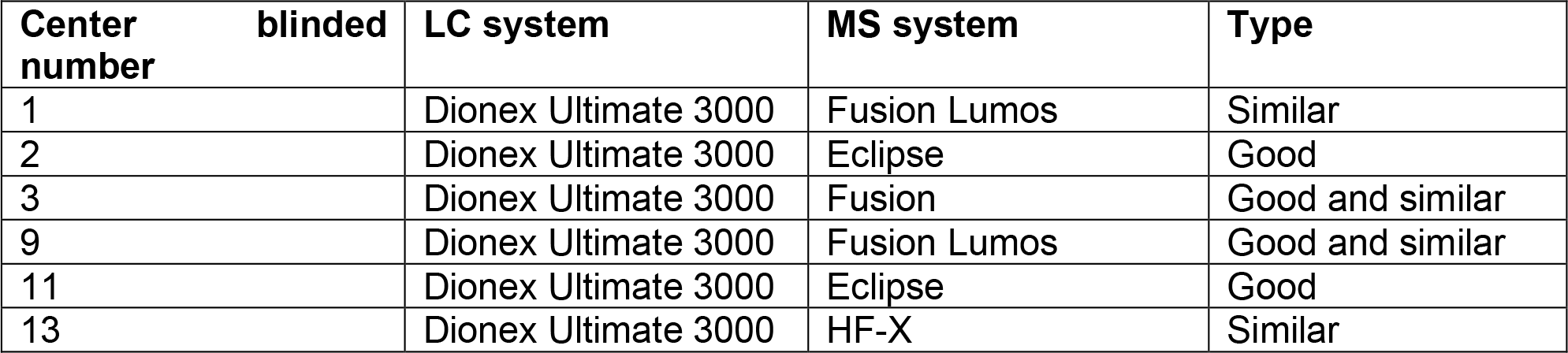
The specific LC and MS systems employed by each of the six core facilities involved in our study.

We consolidated the data from 6 core facilities as detailed in **Supplementary Data 1**. As depicted in **Fig. 2a-b**, even when selecting core facilities that outperformed others based on previously mentioned criteria, the overlap of quantified proteins in identical samples across different centers was only about 11%. In contrast, for core facilities using similar instrumentation, this overlap was 18%. When considering proteins quantified consistently across all three replicates, the overlap percentages were 8% and 14% for good and similar centers, respectively. These findings highlight the impact of instrumentation on the variability of protein corona analysis outcomes. Moreover, protein intensities between different core facilities did not exhibit a consistent pattern, regardless of whether the centers were categorized as ‘good’ and/or ‘similar’ (**Fig. 2c**).

**Fig. 2.**
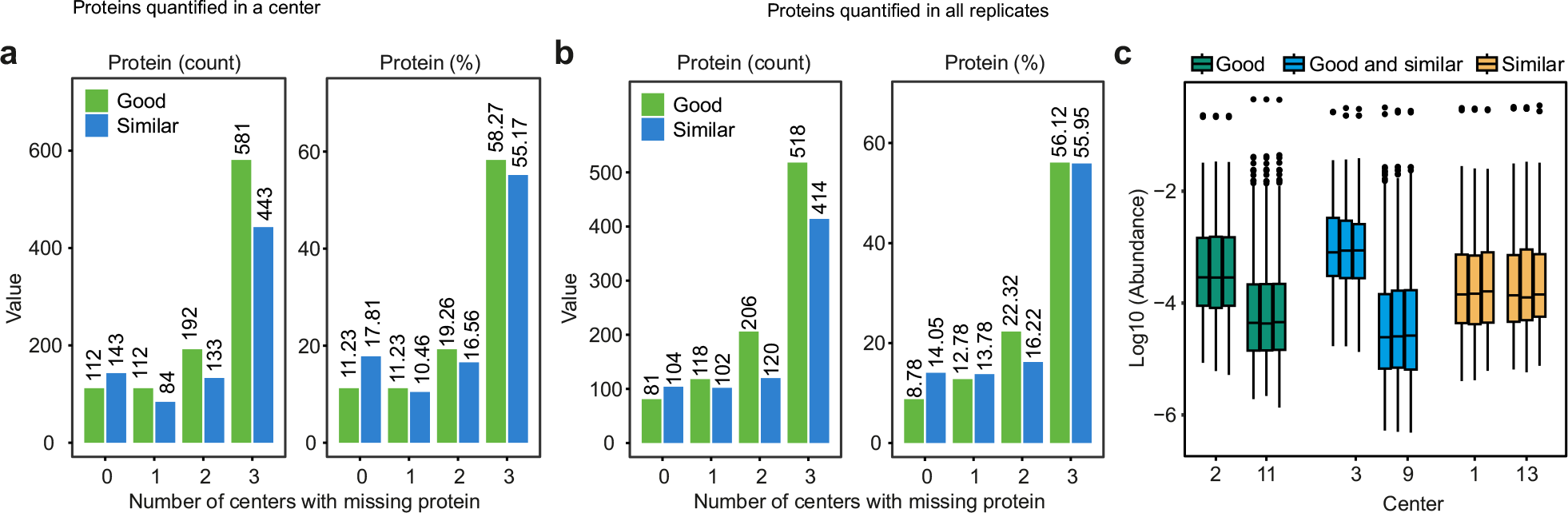
Data overlaps among core facilities reporting better results (good) or having similar instrumentation (similar). **a**, The overlap in the quantified proteins among good or similar centers with regards to protein count and percentage. **b**, The overlap in the quantified proteins among good or similar centers with regards to protein count and percentage, limited to proteins quantified in all replicates. **c**, Distribution of protein-level intensities for the 6 cores (center line, median; box limits contain 50%; upper and lower quartiles, 75 and 25%; maximum, greatest value excluding outliers; minimum, least value excluding outliers; outliers, more than 1.5 times of upper and lower quartiles). All analyses were based on three technical replicates.

The hierarchical clustering in **Fig. 3a-b** shows data completeness captured by each core facility for good and similar centers, respectively. The upset plots shown in **Fig. 3c-d** demonstrate the uniqueness of the data obtained from the different core facilities for good and similar centers, respectively. These plots were made with all the proteins quantified across all the cores and demonstrate that difference (uniqueness) of the data from each core still outweighs their similarities, despite streamlining sample preparation and the instrumentation used. Note that similarities are higher for centers using similar instruments.

**Fig. 3.**
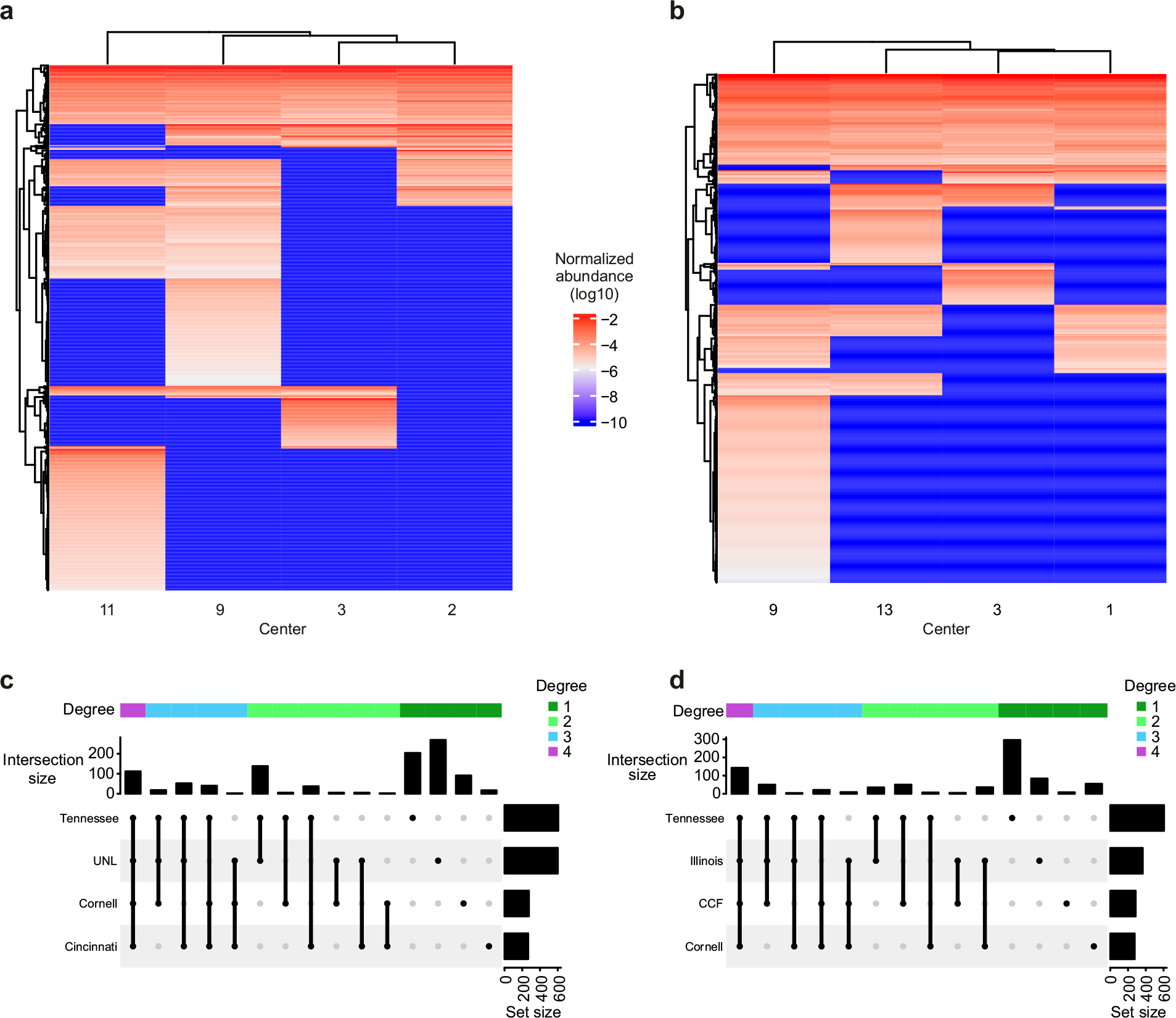
Uniqueness of the data obtained from the different core facilities. **a-b**, Hierarchical clustering of quantified proteins across the 4 good core facilities and 4 core facilities with similar instrumentation, respectively. **c-d**, Upset plot showing the variabilities in the number of detected proteins across the 4 good core facilities and 4 core facilities with similar instrumentation, respectively. All analyses were based on three technical replicates.

### The impact of unifying sample preparation protocol, instrumentation, database search and data processing

As a final step, we subjected the raw data from the 4 centers with similar instrumentation to a unified database search. We have already shown that on top of the sample preparation protocol and the instruments used during sample analysis, database search adds a new layer of variability. These variabilities can be introduced by using different search settings such as false-discovery rates (FDR), inclusion of different fixed and variable post-translational modifications, the number of missed cleavages and the sequence database used in the search.

We included Cysteine carbamidomethylation as a variable modification, since some centers had not specified if the reduction and alkylation of proteins had been performed. Carbamidomethylated peptides were used in the quantification of proteins as well. As for other variable modifications, we only included the default methionine oxidation and acetylation of protein N-termini. Similar to previous studies by us^15^ and others^25^, we included a 1% FDR at both the protein and peptide levels. We only searched for specific tryptic peptides and allowed up to two missed cleavages which is routine procedure. We only applied the parameters that are well-accepted in the community^26^, however, it should be noted that this uniform search does not undermine the validity of the previous database searches performed by the core facilities individually.

In the uniform search, we quantified 370 proteins (**Supplementary Data 2**), which is significantly lower than the combined number of proteins (n=1824) individually reported by the centers. A centralized FDR control, and the application of the identical database search and software is expected to reduce the number of quantified proteins in the aggregated search, as we have also reported.^15^

As shown in **Fig. 4a**, the application of the uniform search substantially improved the number and percentage of the shared proteins in the data retrieved from the 4 core facilities using similar instrumentation. However, due to the reduction of the number of quantified proteins in the uniform database search, the percentage of shared proteins is a more accurate way of comparing performance between the individual searches and the uniform search, than the number of quantified proteins. For proteins with no missing values across centers using similar instruments, the percentage of shared proteins increased from 18% in the individual searches to 40% in the uniform search. These results are consistent with our previous study, where using a uniform database search, we found an overlap of 35.3% among the top 5 facilities (among 15 centers) ^15^. The hierarchical clustering in **Fig. 4b** also shows that the data completeness across different centers was dramatically improved (compared to **Fig. 3b**). Overall, these results show that database search has a higher impact on the data homogeneity than the other parameters that were investigated in this study.

**Fig. 4.**
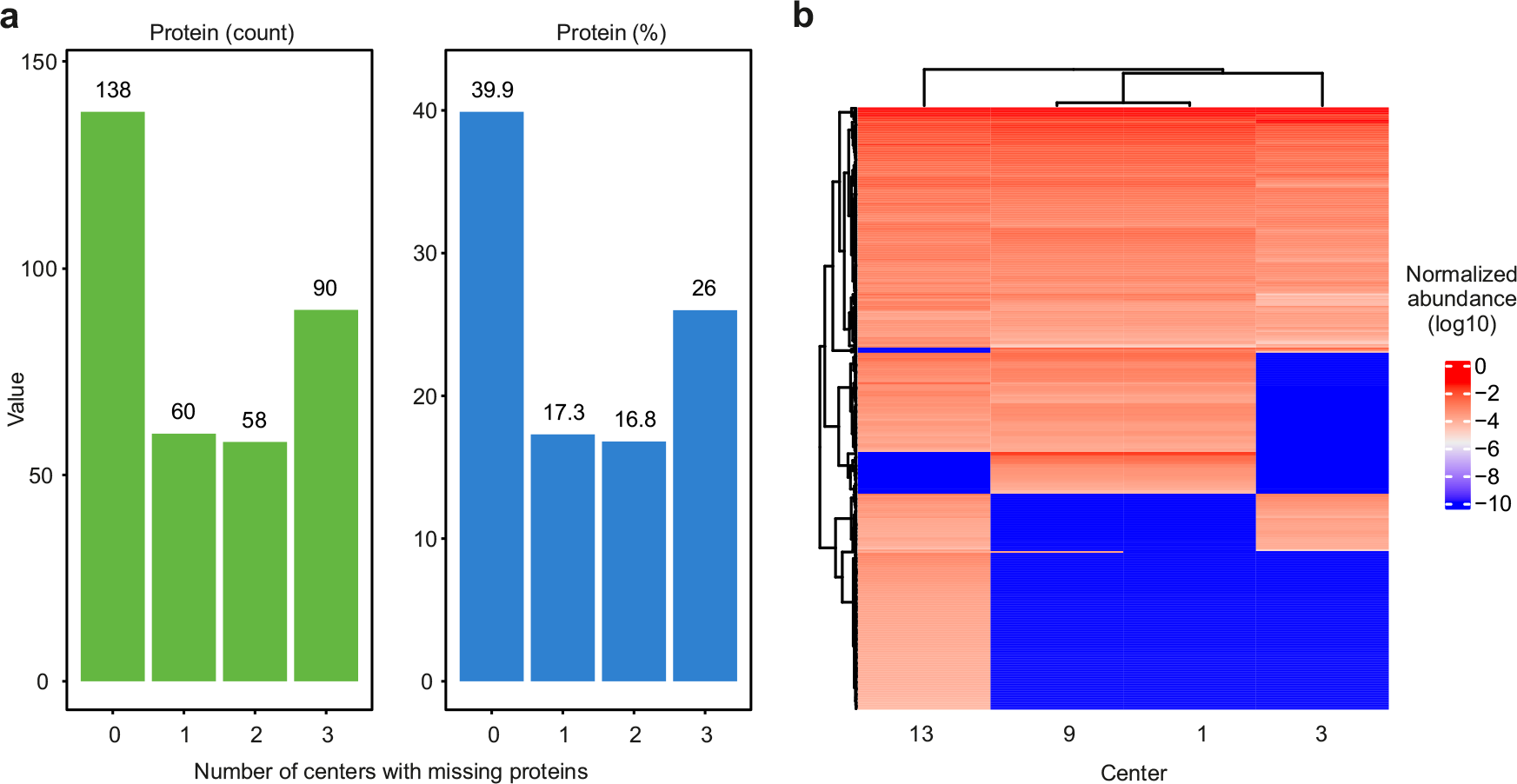
A uniform database search dramatically improves data homogeneity across different core facilities. **a**, The number and percentage of shared proteins with or without missing values in the uniform database search of LC-MS/MS data of the 4 centers using similar instruments, respectively. **b**, Proteins detected in the uniform database search of LC-MS/MS data of the 4 centers using similar instruments. All analyses were based on three technical replicates.

## Discussion

The protein corona spontaneously forms on nanoparticles when exposed to biological tissues and fluids. Recognizing its critical role in influencing the efficacy and safety of nanotechnologies and nanomedicines, extensive research has been dedicated to characterizing the protein corona composition in terms of protein identity and abundance.^14^ Despite numerous studies documented in the literature, efforts to reconcile data from independent studies and consolidate protein corona datasets for predicting nanoparticle’ pharmacokinetics and biological fates are still limited.^27^ While significant progress has been made in standardizing essential characterization methods for nanomedicines to ensure reproducibility and robustness across different research centers^28^, a standardized protocol for analyzing the protein corona composition remains absent. This gap highlights a critical need in the field of nanomedicine, as the protein corona plays a pivotal role in determining the biological identity and behavior of nanoparticles.^6^

MS-based proteomics is the preferred method for characterizing the protein corona. While LC-MS typically offers robust and reproducible data on cell and tissue samples within the same experimental framework^29^, its application to protein corona and plasma-related samples often encounters challenges that restrict proteome coverage across different studies. A significant challenge is the broad dynamic range of protein concentrations in plasma. For example, albumin alone constitutes approximately 55% of the total protein mass in plasma. This dominance by a few high-abundance proteins can mask the presence of lower-abundance proteins, which are crucial for comprehensive proteomic analysis and accurate characterization of the protein corona.^30^ In fact, seven most abundant plasma proteins comprise 85% of total protein in plasma^30^ and upon digestion of plasma proteome, peptides from such abundant proteins crowd the mass spectra, making it challenging to quantify the other present proteins with lower abundance. This issue has been partially mitigated in the past by several depletion strategies that exploit immunodepletion spin columns, immunodepletion-LC, magnetic beads, and even nanoparticlesthemselves.^10, 31, 32^ Such strategies are used to deplete albumin and other abundant proteins before sample analysis. We have recently also introduced a novel methodology where spiking a fine-tuned concentration of phosphatidylcholine and a single nanoparticle were shown to deplete the 4 most abundant proteins in plasma, reducing their cumulative representation (MS intensities) from 90% to under 17% in the whole plasma^33^ and enhancing the plasma proteome coverage by 446% (from 322 to 1436 plasma proteins). Despite these challenges, thousands of proteins have been reliably quantified in plasma in landmark studies, leading to the discovery of distinct disease-based biomarkers.^34, 35, 36^

Altogether, variations in proteome coverage introduce bias in data interpretation by neglecting low abundant proteins that could otherwise be genuine targets or disease biomarkers. The need for comprehensive coverage of the proteome is felt even more in the analysis of nanoparticle protein corona, since the presence and absence of a biomarker is even more important.

Previously we showed that there are significant variations in proteome coverage of identical protein corona samples across 17 core facilities. However, there was a good agreement in LC-MS data among the shared proteins across different core facilities, showing that one of the main challenges is achieving a high proteome coverage^13^. We showed that these variations arise from using different sample preparation protocols, various LC-MS workflows and instruments, as well as database search parameters and data processing^13^. In a follow-up study, we showed that implementing a uniform database search and data processing pipeline, can drastically contribute to data homogeneity (i.e., from 1.8% to 16.2% among top 11 facilities).^15^

In this study, we demonstrated that standardizing the sample preparation protocol, instrumentation, database search, and data processing significantly increases the percentage of shared proteins identified across different facilities in a stepwise manner. Our findings confirm that establishing standard protocols for LC-MS analysis of protein corona can greatly enhance the consistency of proteomics data from various centers. Without such standardized protocols, it is challenging, if not impossible, to expect different core facilities to adopt identical settings for each step of the process. This difficulty arises from the diversity of available analytical tools, such as various types of analytical columns and database search engines. Hypothetically, homogeneity can be further improved by unifying all the instrumental settings, for example including the duration of the gradient, fragmentation techniques, number of scans, resolution and so on. Standardizing these elements is crucial for improving reproducibility and reliability in nanoparticle protein corona research but will remain a formidable challenge.

In summary, the results of this study demonstrated the significant impact of unified protocols on reducing heterogeneity in identical protein corona characterization across different proteomics facilities. By implementing standardized sample preparation, using consistent instrumentation, and harmonizing data processing parameters, we achieved a significant improvement in the homogeneity of the final protein corona outcomes. This highlights the critical role of standardized practices in enhancing data reproducibility and also emphasizes the potential barriers in their implementation due to the diverse capabilities and resources of different laboratories. Despite these challenges, our approach illustrates the possibility of significant improvements in data quality and consistency, paving the way for achieving more reliable protein corona analysis.

Moving forward, the establishment of universal standards for the analysis of the protein corona will be crucial for the advancement of nanomedicine, ensuring that findings from different studies are comparable and that the biological implications of nanoparticle-protein interactions are understood with greater clarity. The adoption of such standards promises to bridge the gap between nanoparticle research and clinical application, ultimately enhancing the development of nanomedicine-based diagnostics and therapeutics.

## Materials and Methods

### Materials

Healthy human plasma protein was obtained from Innovative Research (www.innov-research.com) and diluted to a final concentration of 55% using phosphate buffer solution (PBS, 1X). Plain polystyrene nanoparticles (∼ 80 nm) were provided by Polysciences. (www.polysciences.com).

### Protein corona formation on the surface of nanoparticles

For protein corona formation, nanoparticles were incubated with 55% plasma (with nanoparticles’ concentration of 0.2 mg/ml) for 1h at 37 °C at a constant agitation (total volume: 9’
s1.5 mL Eppendorf tubes). To remove unbound and plasma proteins only loosely attached to the surface of nanoparticles, protein-nanoparticles complexes were then centrifuged at 14,000 ’ g for 20 minutes, the collected nanoparticles’ pellets were washed twice more with cold PBS under the same conditions, and the final pellet was collected for preparation for LC-MS analysis.

### LC-MS/MS sample preparation

The protein corona pellets (i.e., 6 separate batches) obtained in the previous step were mixed with 25 μL buffer containing 50 mM ammonium bicarbonate (AMBIC) and 1.6 M Urea. Then 2.5 μL of dithiothreitol (DTT, Pierce BondBreaker, neutral pH) was added to 10 mM final concentration and incubated at 37 oC for 60 minutes using a Thermomixer. Samples were then cooled down to room temp, vortexed, and spun down. 2.7 μL of freshly made iodoacetamide (IAA) was then added to a 50 mM final concentration and samples incubated for 1 h at room temperature in dark. 3 μL of DTT was added to a 10 mM final concentration and incubated at room temperature for 15 minutes to quench the IAA. Then freshly diluted trypsin dissolved in protein digestion buffer (from Promega), was added to the samples at a ratio of 1:50 of trypsin:protein and incubated overnight at 37 oC in a Thermomixer. The following day, samples were removed from Thermomixer and cooled down to room temperature. Samples were then centrifuged at 16,000 ’s g for 20 minutes to remove the nanoparticles from supernatant. All supernatants (peptides) were transferred to separate new low binding Eppendorf tubes followed by adding 5% formic acid (FA) to the samples to adjust the pH between 2 and 3. The peptides were then desalted and cleaned using c18 StageTips in new low binding Eppendorf tubes. It is noteworthy that the c18 StageTips were first initiated using 80% acetonitrile (ACN) and 5% formic acid (FA) and equilibrated with 5% FA followed by loading and washing the sample with 5% FA. The peptides were eluted sequentially first using 30% ACN and 5% FA solution, followed by 80% ACN and 5% FA. Finally, all 6 identical eluates of prepared peptides were mixed, shaken, and aliquoted again to 6 identical batches in new low binding Eppendorf tubes and dried in a speed vacuum centrifuge. The dried peptides were then shipped overnight (using FEDEX) in dry ice with guaranteed next-morning delivery. It is noteworthy that safe arrival of all samples was confirmed by each core facility on the next day.

### Characterization

DLS and zeta potential analyses were performed to measure the size distribution and surface charge of the nanoparticles before and after protein corona formation using a Zetasizer nano series DLS instrument (Malvern company). A Helium Neon laser with a wavelength of 632 nm was used for size distribution measurement at room temperature. TEM was carried out using a JEM-2200FS (JEOL Ltd.) operated at 200kV. The instrument was equipped with an in-column energy filter and an Oxford X-ray energy dispersive spectroscopy (EDS) system. 20 μl of the bare nanoparticles were deposited onto a copper grid and used for imaging. For protein corona–coated nanoparticles, 20 μL of sample was negatively stained using 20 μl uranyl acetate 1%, washed with DI water, deposited onto a copper grid, and used for imaging. Protein corona composition was also determined using LC-MS/MS. LC-MS/MS analyses were carried out at 6 different proteomics cores across the United States that performed better based on our previous findings and had similar instrumentation (i.e., LC and mass types).

### LC-MS/MS analysis and proteomics data processing

Details of sample preparation, LC-MS/MS analysis, and data processing protocols are included in **Supplementary Information** files for each core. To avoid any possible bias in the final protein corona outcomes, we asked all cores to resuspend the dried peptides in 30 μL of 0.1%FA and 2% ACN (in water), split it to 3 parts and inject ∼ 5 μL of each for 120 min analysis in the LFQ mode. Moreover, to avoid any bias in instrumentation, search parameters and raw data processing, we asked the 4 cores with similar instrumentation to consider M oxidation, acetylation of protein N-termini, cysteine carbamidomethylation, and a maximum of 2 missed cleavages when analyzing the data. In addition, we also asked the 4 cores to analyze the data by MaxQuant and if not possible by proteome discoverer.

## Data availability

Due to the blinding of core names in the current study, and since the .raw files can be traced, the .raw data and associated individual data files are available upon request from corresponding authors. The extracted protein abundances data and relevant outputs of data analysis are provided in supplementary data files cited in the text. **Supplementary Data 1** was used to generate **Figs. 2-3** and **Supplementary Data 1** for **Fig 4** (**Supplementary Data 1-2** as the Source Data).

## Data analysis

First, for each core, data were normalized by total protein intensity in each technical replicate. CVs were calculated based on normalized intensities between technical replicates for each protein. To unify protein IDs, for some cores, the various protein IDs used were converted to UniProt IDs. The data among the cores were combined by UniProt IDs.

## Author contributions

Conceptualization, A.A.A., A.A.S. and M.M.; project organization, resources and funding acquisition, M.M. and A.A.S.; methodology and experiment design, M.M., A.A.S. and A.A.A.; experiments and characterizations, A.A.A.; data analysis and visualization, H.G., A.A.S., S.M.M. and A.A.A.; writing—original draft, A.A.A., A.A.S., H.G. and M.M.; writing, review & editing, all co-authors.

## Ethics declarations Competing interests

Morteza Mahmoudi discloses that (i) he is a co-founder and director of the Academic Parity Movement (www.paritymovement.org), a non-profit organization dedicated to addressing academic discrimination, violence and incivility; (ii) he is a co-founder of Targets’ Tip; and (iii) he receives royalties/honoraria for his published books, plenary lectures, and licensed patents.

## Acknowledgement

We acknowledge all 6 core facilities for their efforts and contributions to the current study.

