## Supplementary material for "Standardizing protein corona characterization in nanomedicine: a multi-center study to enhance reproducibility and data homogeneity": Supp Information

**Supplementary Fig. 1. Characterizations of the pristine and protein corona-coated nanoparticles. a,** and **b**, DLS analysis of pristine and protein corona–coated nanoparticles (respectively) and the corresponding replicates. Representative TEM images of nanoparticles before (**c**, and **d**) and after (**e**) formation of protein corona The table presents the summarized average size, surface charge and polydispersity index (PDI) of pristine and protein corona-coated nanoparticles, as representative. (PC: protein corona)

**Supplementary materials and methods**

**BCA analysis**

To estimate the concentration of the proteins in identical batches of protein corona-coated nanoparticles, we measured the protein concentration using bicinchoninic acid assay (BCA). BCA analysis confirmed the consistency of protein amount in the identical batches of protein corona coated nanoparticles. In the low concentrations of BSA, where there is a meaningful linear relationship between the concentrations and absorption values, the protein concentration was calculated about ~ 1.6 µg in each batch after protein corona formation. In general, there is 20-40% loss in protein concentration during digestion and sample preparations for LC-MS analysis and therefore, we estimate that each vial contains less than 1 µg protein. Since we asked all proteomics core to resuspend and split the sample into three technical replicates, and therefore, in each individual analysis ~ 350 ng protein was analyzed by LC-MS system. It is noteworthy that none of the cores reported any issues such as column clogging during the measurements.

**LC-MS**

The following sections describe the experimental details and instrumentation of LC-MS/MS analysis provided by each individual proteomic core facility and reported here as received (with some minor changes for consistency of units, etc.). The 6 different proteomics core facilities that contributed to this study include Cornell University ([Link](https://www.biotech.cornell.edu/core-facilities-brc/facilities/proteomics-metabolomics-facility)), Cleveland Clinic Lerner Research Institute ([Link](https://lerner.ccf.org/cores/proteomics-metabolomics/)), University of Cincinnati ([Link](https://www.artsci.uc.edu/departments/chemistry/core-facilities/mass-spectometry-facility/mass-spectometry-facility-staff.html)), University of Tennessee ([Link](https://www.uthsc.edu/research/institutional-cores/pmc/)), University of Nebraska–Lincoln ([Link](https://biotech.unl.edu/proteomics-and-metabolomics#tab6)), University of Illinois ([Link](https://biotech.illinois.edu/proteinsciences)). The centers were blindly numbered as 1, 2, 3, 9, 11, and 13 in random order corresponding to our previous report.^1^

**Center #1:**

The digest was analyzed in triplicate by capillary column LC-tandem MS using a LC gradient from 2 to 48% acetonitrile in 120 minutes. The CID spectra was searched against the human SwissProtKB database. Five μL volumes of the extract were injected and the peptides eluted from the column by an acetonitrile/0.1% formic acid gradient at a flow rate of 0.3 μL/min were introduced into the source of the mass spectrometer online. The micro-electrospray ion source was operated at 2.5 kV. The digest was analyzed using the data dependent multitask capability of the instrument acquiring full scan mass spectra to determine peptide molecular weights and product ion spectra to determine amino acid sequence in successive instrument scans. The data were searched against the human SwissProtKB protein database with the program Sequest and MSFragger. Protein and peptide validations were performed with the program Scaffold to <1% FDR at the protein and peptide levels.

The LC-MS system was a Dionex Ultimate 3000 nano-flow HPLC interfacing with a ThermoScientific Fusion Lumos mass spectrometer system. The HPLC system used an Acclaim PepMap 100 precolum (75 μm x 2 cm, C18, 3 μm, 100 A) followed by an Acclaim PepMap RSLC analytical column (75 μm x 15 cm, C18, 2 μm, 100 A).

**Center #2:**

The dried sample was resuspended in 30 µL of 0.1% formic acid and 5 µL of the sample was analyzed in triplicate by nanoLC-MS/MS (Dionex Ultimate 3000 nano-flow HPLC connected to Orbitrap Eclipse) and was searched against a combined database containing common contaminants and the *Homo sapiens* (UP000005640) database using Proteome discoverer version 2.4 with the Sequest HT search algorithm (Thermo scientific) using a multiconsensus by sample run Label Free Quantitation (LFQ) workflow which provides abundance values for each sample with ratios and still shows the Sequest scores, peptides and PSMs for each file individually.

**Center #3:**

The tryptic digest sample was reconstituted in 30 μL of 2% acetonitrile (ACN) with 0.5% formic acid (FA) for nanoLC-ESI-MS/MS analysis and aliquoted to three separate sample vials. The analysis was carried out using a Dionex Ultimate 3000 nano-flow HPLC connected to an Orbitrap FusionTM TribridTM (Thermo-Fisher Scientific, San Jose, CA) mass spectrometer equipped with a nanospray Flex Ion Source, and coupled with a Dionex UltiMate 3000 RSLCnano system (Thermo, Sunnyvale, CA).^2, 3^ The peptide samples (5 μL) of each aliquot were injected onto a PepMap C-18 RP nano trapping column (5 µm, 100 µm i.d. x 20 mm) at 20 µL/min flow rate for rapid sample loading and then separated on a PepMap C-18 RP nano column (2 µm, 75 µm i.d. x 25 cm) at 35°C. The tryptic peptides were eluted in a 120-min gradient of 5% to 35% ACN in 0.1% formic acid at 300 nL/min, followed by a 7-min ramping to 90% ACN-0.1% FA and an 8-min hold at 90% ACN-0.1% FA. The column was re-equilibrated with 0.1% FA for 25 min prior to the next run. The Orbitrap Fusion was operated in positive ion mode with spray voltage set at 1.1 kV and source temperature at 275°C. External calibration for FT, IT and quadrupole mass analyzers were performed. In data-dependent acquisition (DDA) analysis, the instrument was operated using FT mass analyzer in MS scan to select precursor ions followed by 3 second “Top Speed” data-dependent CID ion trap MS/MS scans at 1.6 m/z quadrupole isolation for precursor peptides with multiple charged ions above a threshold ion count of 10,000 and normalized collision energy of 30%. MS survey scans at a resolving power of 120,000 (FWHM at m/z 200), for the mass range of m/z 375-1575. Dynamic exclusion parameters were set at 50 s of exclusion duration with ±10 ppm exclusion mass width. All data were acquired under Xcalibur 4.4 operation software (Thermo-Fisher Scientific).

The DDA raw files with MS and MS/MS were subjected to database searches using MaxQuant version 2.3.1.0 in an LFQ workflow against *Homo sapiens* Uniprot database containing 26,019 sequences along with a regular contaminant (244 entries) database.  Oxidation on M, acetylation of protein N-termini, and deamidation on N were specified as dynamic modifications, while carbamidomethylation on C was specified as a static modification, and a maximum of 2 missed cleavages by trypsin digestion was allowed.  The peptide mass tolerance in MS mode was set to 10 ppm and MS/MS tolerance was set to 0.6 Da. The estimated false discovery rate (FDR) thresholds for protein, peptide and modification site were specified at a maximum of 1%. The minimum peptide length was set at 6 and unique and razor peptide intensities were used. All the other parameters in MaxQuant were set to default values.^4^

Relative quantitation of identified proteins between the three technical replicates was determined by the LFQ workflow in MaxQuant. The precursor abundance intensity (the output of the MaxLFQ algorithm,^5^ for each peptide identified by MS/MS in each replicate was automatically determined and their unique plus razor peptides for each protein in each replicate were summed and used for calculating the protein abundance.

**Center #9:**

The submitted desalted/dried peptide sample (~1.6 µg) was re-dissolved in 30 µL of loading buffer (3% acetonitrile, 0.1% TFA), and 5 µL (~0.267 µg) was analyzed in triplicate (technical replicates) using LC-MS-MS method with 160 min LC gradient for peptide/protein identification and LFQ. The following instrumentation was used:

HPLC: Ultimate 3000RSLCnano, Thermo Fisher

Trap column: Acclaim PepMap 100, 75 µm x 20 mm, C18, 3 µm, 100Å, Thermo Fisher

Injection volume/mode: 5 µL/µLPickUp

Loading Buffer: 3% acetonitrile with 0.1% TFA

Loading flow rate and duration: 5 µL/min for 5 min

Column: Acclaim PepMap RSLC, 75 µm x 500 mm (ID x Length), C-18, 2 µm, 100 Å, Thermo Fisher

Solvent A: 0.1% formic acid in water, LC/MS grade, Thermo Fisher

Solvent B: 0.1% formic acid in acetonitrile, LC/MS grade, Thermo Fisher

LC Flow rate: 300nl/min

Column temperature: 40 °C

Gradient: 0min-3%B, 4min-3%B, 5min-5%B, 110min-23%B, 120min-30%B, 123min-90%B,

133min-90%B, 136min-3%B, 160min-3%B

MS: Orbitrap Fusion Lumos, Thermo Fisher

Data dependent analysis (DDA): 3sec cycles

MS scan (full): Analyzer - Orbitrap, resolution-120,000 (FWHM, at m/z=200)

Scan Filters:	MIPS mode - Peptide

Intensity ≥ 10,000

Charge state - 2-6

Dynamic exclusion – 45 sec

MS2 scan (full): Quadrupole isolation window - 0.7 m/z,

Activation - HCD (30%)

Analyzer - Orbitrap, Resolution 30,000 (FWHM, at m/z=200)

Post-Acquisition Analysis of raw MS data

Proteome Discoverer 2.4, Thermo Fisher

Peptide/Protein Identification

Search engine: Sequest HT

Target Database: SwissProt, TaxID 9606 (Homo sapiens), v.2022-10-22, 42315 entries

Decoy Database: reversed target database

Enzyme: Trypsin (full)

Dynamic modification: Oxidation of Met, Met loss, acetylation of the protein N-terminus

Static modification: Carbamidomethylation of Cys

Precursor and fragment ion mass tolerance: 10ppm and 0.02Da, respectively

Validation and filtering at PSM level (q value): Percolator, FDR ≤0.01

Validation and filtering at peptide level (q value): Qvality algorithm, FDR ≤0.01

Identification of protein or protein group: At least one validated peptide sequence

unique to a protein or a protein group

Protein groups: Strict parsimony principle applied

Validation at protein level (Experimental q value): strict - FDR≤0.01, relaxed - FDR≤0.05

Feature Detection

Min Trace Length: 5

Min # Isotopes: 2

Max ΔRT of Isotope Pattern Multiplets: 0.2min

Chromatographic Alignment

Max RT shift: 5min

Mass tolerance: 10ppm

Feature Linking/Mapping

RT tolerance: 0 (automatic)

Mass tolerance: 0 (automatic)

Min S/N threshold: 5

Peptide/Protein Quantification:

Quantification: LFQ - (Precursor Ion Area Detection)

Peptides to use: Unique + Razor

Peptide uniqueness: Protein Group

Peptide Abundance: MS Peak Area

Normalization mode: Total Peptide Amount

Protein abundance: Summed abundances of assigned peptides

Peptide group abundance: Mean of bio-replicate abundances

Protein group abundance: Mean of bio-replicate abundances

Ratio calculation: Based on summed abundances

Hypothesis test: t Test (Individual Proteins)

Adjusted p-value: Benjamini-Hochberg method.

**Center #11**

5 μL of each digest was injected in triplicate and run by Dionex Ultimate 3000RSLCnano susing a 2h gradient on a Waters CSH 0.075 mmx250 mm C18 column feeding into a Thermo Eclipse mass spectrometer run in OT-IT mode, using HCD for fragmentation. The quantitation of the proteins was performed using Proteome Discoverer (Thermofisher; version 2.4). All MS/MS samples were searched using Sequest set up to search the cRAP_20150130.fasta (125 entries); uniprot-human_20221103 database (80581 entries) assuming the digestion enzyme trypsin. Sequest was searched with a fragment ion mass tolerance of 0.6 Da and a parent ion tolerance of 15.0 PPM. Carbamidomethylation of cysteine, deamidated of asparagine and glutamine, and oxidation of methionine were specified as variable modifications. Peptides were validated by Percolator with a 0.01 posterior error probability (PEP) threshold. The data were searched using a decoy database to set the FDR to 1% (high confidence). The peptides were quantified using the precursor abundance based on intensity. The peak abundance was normalized using total peptide amount. Normalized, scaled and raw abundances are reported. Only proteins and peptides identified and quantified in at least one of the technical replicates were reported. Protein with number of peptides and PSM as low as 1 were reported here because of the overall low protein coverage.

**Center #13:**

The peptide sample was suspended in a solution of 0.1% FA in 2% ACN, and 300 ng from each sample was injected into an UltiMate 3000 RSLCnano system. The peptides were separated using a 25 cm Acclaim PepMap 100 C18 column (Thermo Scientific) and mobile phases of 0.1% FA (A) and 0.1% FA in 80% ACN (B) at a flow rate of 300 nL/min. The gradient started at 5% B, increased to 35% B over 40 minutes, and then increased to 50% B over 5 more min; this was followed by column washing and equilibration. The column was maintained at 50°C over the course of the run.

The peptides were analyzed with a Q Exactive HF-X mass spectrometer (Thermo Scientific) in the positive mode. MS1 scans from 350 to 1500 m/z were acquired at 120k resolution (3e6 AGC; 60 ms max IT), followed by HCD fragmentation (30 NCE) of the 15 most abundant ions. MS2 scans were acquired at 30k resolution with an isolation window of 1.0 m/z and a dynamic exclusion time of 20 s (5e4 AGC; 35 ms max IT).

The raw LC-MS data was searched against the *Homo sapiens* Uniprot reference proteome (March 2023) with MaxQuant v2.0.1.0. Trypsin was specified as the enzyme with a maximum of 2 missed cleavages; minimum peptide length was set to 6. Variable modifications of methionine oxidation and N-terminal acetylation were also added to the search. The precursor mass tolerance was 20 ppm for the first search and 10 ppm for the main search, while the fragment mass tolerance was set to 20 ppm. The match between runs function was enabled with a match time window of 0.7 min, and the FDR was set to 1% at the PSM and protein levels. LFQ was also performed with MaxQuant using a minimum ratio count of two.

**Acknowledgements:**

We thank the Proteomics and Metabolomics Facility of Cornell University for providing the mass spectrometry data and NIH SIG grant 1S10 OD017992-01 support for the Orbitrap Fusion mass spectrometer. This work was supported by the Vincent Coates Foundation Mass Spectrometry Laboratory, Stanford University Mass Spectrometry (RRID:SCR_017801). This work was supported in part by NIH P30 CA124435 utilizing the Stanford Cancer Institute Proteomics/Mass Spectrometry Shared Resource.
